# The tau oligomer antibody APNmAb005 detects early-stage pathological tau enriched at synapses and rescues neuronal loss in long-term treatments

**DOI:** 10.1101/2022.06.24.497452

**Authors:** Hwan-Ching Tai, Haou-Tzong Ma, Shien-Chieh Huang, Meng-Fang Wu, Chia-Lin Wu, Yen-Ting Lai, Zhong-Lin Li, Richard Margolin, Anthony J. Intorcia, Geidy E. Serrano, Thomas G. Beach, Madhaven Nallani, Bradford Navia, Ming-Kuei Jang, Chin-Yin Tai

## Abstract

Numerous tau immunotherapies are being developed against Alzheimer’s disease (AD), but it has been challenging to specifically target early-stage tau aggregates using conformation-dependent antibodies. Here, we report a monoclonal antibody, APNmAb005, that recognized a conformational epitope associated with tau oligomers. In AD brain extracts, mAb005 preferentially recognized oligomeric tau in the synapse over monomeric tau in the cytosol. In the prefrontal cortex and hippocampus, mAb005 immunoreactivity was strongly present in early-stage AD but surprisingly diminished in late-stage AD (Braak stage VI). mAb005 also recognized aggregates in 3R tauopathies (Pick’s disease) and 4R tauopathies (corticobasal degeneration and progressive supranuclear palsy), including those in astrocytes and oligodendrocytes. In rTg4510 mice (P301L tau), mAb005 immunoreactivity first appeared in distal neurites but much later in neuronal somas. Thus, the mAb005 epitope appears to be associated with early-stage oligomers of tau (esoTau) that accumulate around synapses in AD, which is also detectable in both 3R and 4R tauopathies. In cellular uptake models of tauopathy transmission, mAb005 blocked the formation of intracellular inclusions induced by incubation with rTg4510 mouse brain extracts. Long-term treatments with mAb005 in rTg4510 mice partially rescued synaptic and neuronal loss in the hippocampus without promoting overall tau clearance. Our data suggest that immunotherapies targeting esoTau enriched around synaptic sites may alleviate tau toxicity against synapses and neurons, which may be a promising treatment strategy against AD.

**One Sentence Summary:** A tau-conformer antibody recognizing synaptic oligomers and 3R, 4R, and mixed aggregates in humans rescues neuronal loss in mouse tauopathy models.

## INTRODUCTION

The neuropathology of Alzheimer’s disease (AD) is characterized by the misfolding of β-amyloid peptide (Aβ) and hyperphosphorylated tau (pTau) proteins (*1, 2*). Increasing evidence has shown that the critical toxic agents may be soluble oligomers of Aβ and tau instead of insoluble fibrils (*3-6*). Currently, there is no effective therapeutic strategy to slow down AD progression (*7*). Dozens of immunotherapies against Aβ and tau have undergone clinical trials for AD treatment (*8-10*). So far, the only clinically approved immunotherapy is Aducanumab, a conformational antibody against Aβ oligomers, but it has shown limited clinical benefits thus far (*11, 12*).

As extracellular oligomers have become the prime target in Aβ immunotherapy (*13*), the suitable targets for tau immunotherapy remain unclear. Tau proteins are found in many different cellular compartments of the brain (*14-16*), exhibiting various misfolding conformations (*17, 18*) and propagative strains (*19-21*), with a multitude of proteoforms arising from extensive posttranslational modifications (phosphorylation, acetylation, methylation, glycosylation, etc.) (*22-27*). Based on the prion-like propagation hypothesis (*28-32*), tau immunotherapies should ideally target early-stage transmissive species for early intervention without interfering with native tau proteins and their physiological functions. In AD subjects, the seeding capacity of tau oligomers appears before neurofibrillary tangle deposition (*33*). We also observed widespread accumulation of tau oligomers in AD-affected synapses, which have the potential to be propagative seeds (*34, 35*). Interestingly, in the rare cases of non-demented elderly with extensive plaque and tangle pathologies, their synapses show very little tau oligomer deposition (*6, 36*). Therefore, tau oligomers around synaptic sites, rather than insoluble tau fibrils, may play crucial roles in synaptic dysfunction and cognitive decline.

Previously, eleven tau antibodies have undergone clinical trials against AD (*10, 37*), which may be classified into three classes. Firstly, six of them recognize linear epitopes on native tau protein, lacking specificity for pathological species. Their proposed mode of action (MOA) is to block transmission by targeting extracellular tau (eTau) (*38, 39*). Secondly, four of them target various phosphosites, including pS217 (*40*), pT231 (Pinteon PTN001), pS396 (*41*), and pS422 (*42*). Dozens of phosphosites have been identified on tau, but it remains difficult to assess whether a particular phosphosite may promote toxicity, reduce toxicity, or carry out physiological functions under various contexts (*22, 43, 44*). Thirdly, LY3303560 (*45*) is the only conformation-specific antibody being clinically tested (halted after phase 2 trial), which is the humanized version of MC-1, an IgG against fibrillar aggregates (*46*).

Here, we sought to develop a tau immunotherapy that focuses on pathological species and early intervention, by targeting tau oligomers accumulating around synapses. However, oligomers are metastable species which are generally difficult to generate, store, and immunize against. Previously, we found that encapsulation of Aβ into artificial vesicles led to stabilized oligomers that are conformationally distinct from recombinant fibrils (*47, 48*). Here, we encapsulated tau oligomers into artificial vesicles to create the biomimetics of AD-affected synapses. After injecting them into mice for antibody screening, we identified a monoclonal IgG, APNmAb005, that preferentially recognized a conformational epitope associated with synaptic tau oligomers in AD. The staining patterns of mAb005 in postmortem AD tissues and rTg4510 mice (expressing P301L human tau (*49*)) suggested a preference for early-stage oligomers of tau (esoTau). Treatments with mAb005 partially rescued synaptic and neuronal loss in rTg4510 mice without apparent reductions in tau aggregates. Our results suggest that targeting esoTau to reduce neuronal and synaptic toxicity may be a promising strategy for tau immunotherapy.

## RESULTS

### mAb005 recognizes misfolded tau oligomers

To screen for conformer-specific tau antibodies, we prepared biochemical fractions enriched with misfolded pTau aggregates from AD brain tissues, including the synaptosomes and the sarkosyl-insoluble materials from brain lysates. pTau was detected by AT8 (pS202/pS205) immunoreactivity, a common pathological marker (**Fig 1a-1d)**. Brain-derived tau aggregates were used for dot-blot screening of eleven monoclonal antibodies generated against vesicle-encapsulated tau oligomers **(Fig. 1e)**. The first group of antibodies (mAb005, 010, 011, 013, and 032) only recognized pathological tau species in the sarkosyl-insoluble and synaptosomal fractions of AD brains. Because synaptosomes only contain tau oligomers but not fibrils (*34*), these may be classified as oligomer-recognition antibodies. The second group of antibodies (mAb004 and 033) recognized both normal and pathological tau, similar to the general-purpose tau antibody HT7. A third group (mAb008, 020, 025, and 037) showed intermediate binding preferences between the first and second group, showing weaker reactivity against sarkosyl-soluble tau but stronger reactivity against sarkosyl-insoluble tau.

**Figure 1.**
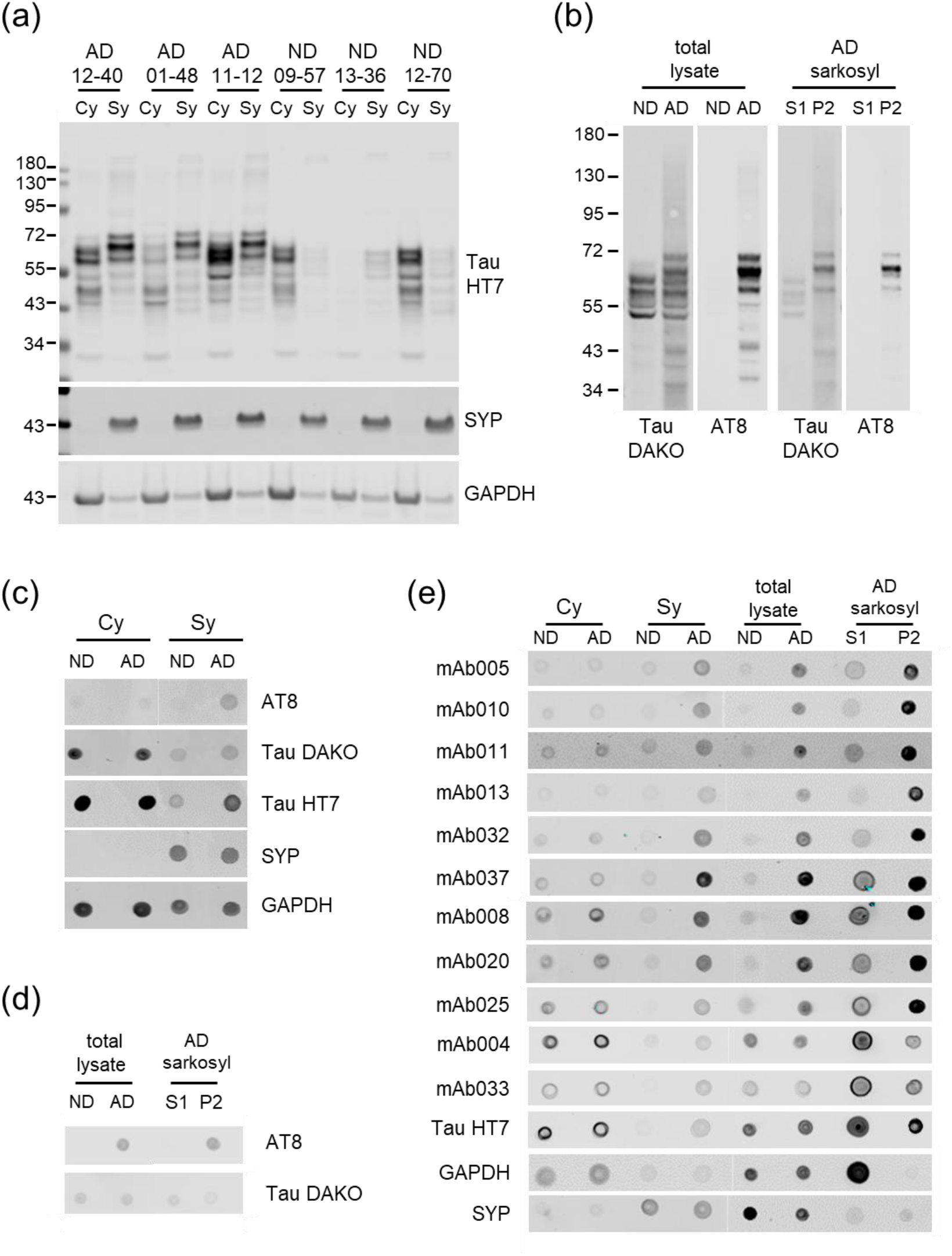
(a) Cytosolic (Cy) and synaptosomal (Sy) extracts from AD and non-demented (ND) subjects, probed against human tau (HT7), synaptophysin (SYP), and GAPDH. (b) AD brain total lysates were separated into sarkosyl soluble (S1) and insoluble (P2) fractions, probed by total tau (DAKO) and pTau (AT8) antibodies. (c) Dot-blot analysis of materials from (a) for pTau, total tau, human tau (HT7), SYP and GAPDH. (d) Dot-blot analysis of materials from (b) for pTau and total tau. (e) Characterization of tau monoclonal antibodies (mAb series) by dot-blotting against materials from (a) and (b).

Four of these antibodies—mAb004, 005, 020, and 037—were further characterized by immunoblotting against human brain materials. mAb005 was distinguished by very low reactivity against tau monomers in the brain lysate of non-demented controls and the sarkosyl-soluble tau of AD subjects. mAb005 was partially reactive toward tau proteins from rTg4510 brain lysates and sarkosyl-insoluble fraction from AD subjects, implying that they retained partial misfolding under denaturing/reducing gel conditions (**Fig. S1**). Among these four antibodies, mAb005 showed the greatest specificity against tau aggregates isolated from AD brains, including high-molecular weight species from size exclusion chromatography (fraction 9 in **Fig. 2a-b**) and high-density species from sucrose gradient centrifugation (30-50% fractions in **Fig. S2**). Against tau aggregates prepared *in vitro*, mAb005 was strongly reactive against oligomers, partially reactive against fibrils, and barely reactive against monomers (**Fig. S2c**). Taken together, our data suggested that mAb005 preferentially recognized a conformational epitope associated with misfolded tau oligomers.

**Figure 2.**
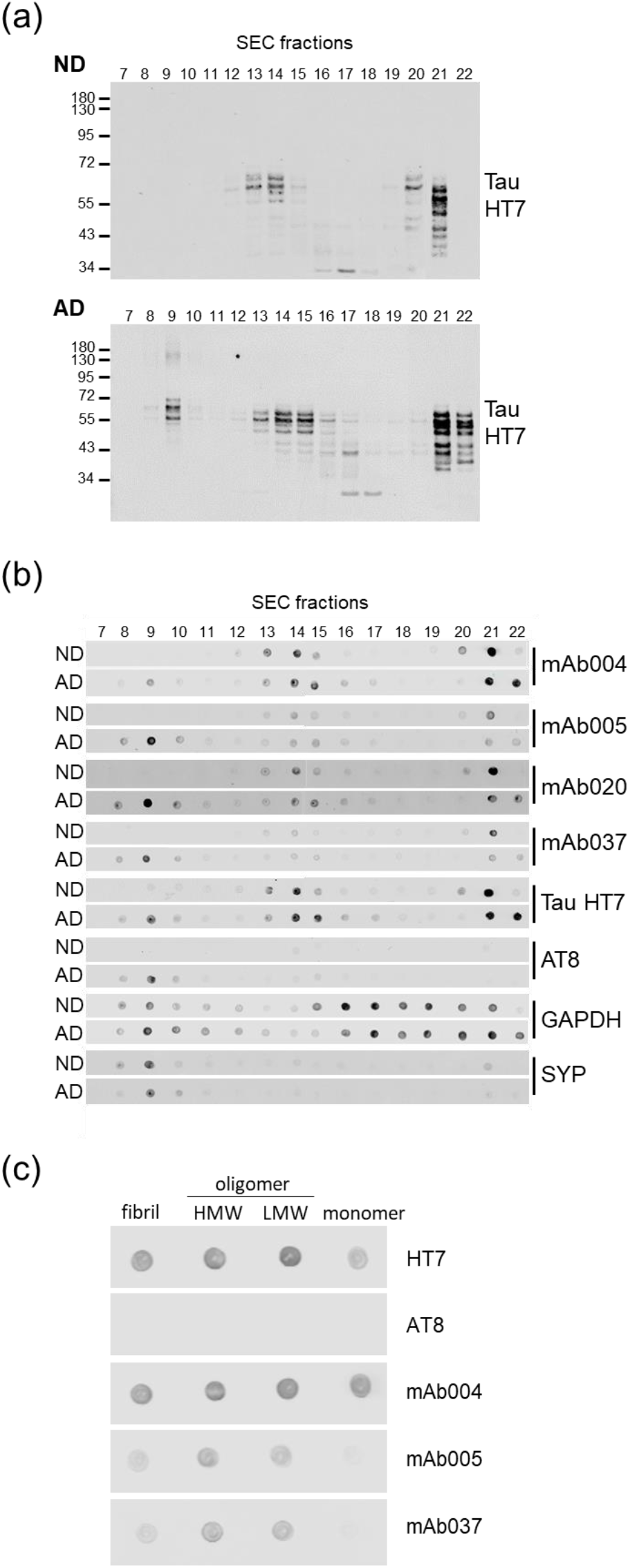
(a) AD brain extracts contain more high-molecular weight tau species (SEC fraction 9) compared to non-demented (ND) controls. Fractions 21-22 contain tau monomers. (b) Dot-blot analysis of SEC fractions using tau monoclonal (mAb series), HT7, AT8 (pTau), GAPDH, and synaptophysin (SYP) antibodies. (c) Dot-blots of tau monoclonal antibodies against recombinant tau aggregates prepared *in vitro*, separated by SEC into fibrils, oligomers (high and low molecular weight), and monomers.

### mAb005-reactive conformers represent early misfolding aggregates

In the hippocampal CA1 and CA3 regions of 3-month-old rTg4510 mice, HT7 immunoreactivity (total human tau) was concentrated in neuronal somas. By contrast, mAb005 immunoreactivity was hardly detectable in somas but concentrated instead in the neuropil layers. In six-month-old rTg4510 mice, HT7 signals became highly prominent in neuronal cell bodies bearing tangles, while mAb005 immunoreactivity had also spread from neuropils into some neuronal somas (**Fig. 3**). This implied that mAb005 recognized an early series of tau misfolding events that originated around distal neurites and perhaps synapses, far away from the cell body.

**Figure 3.**
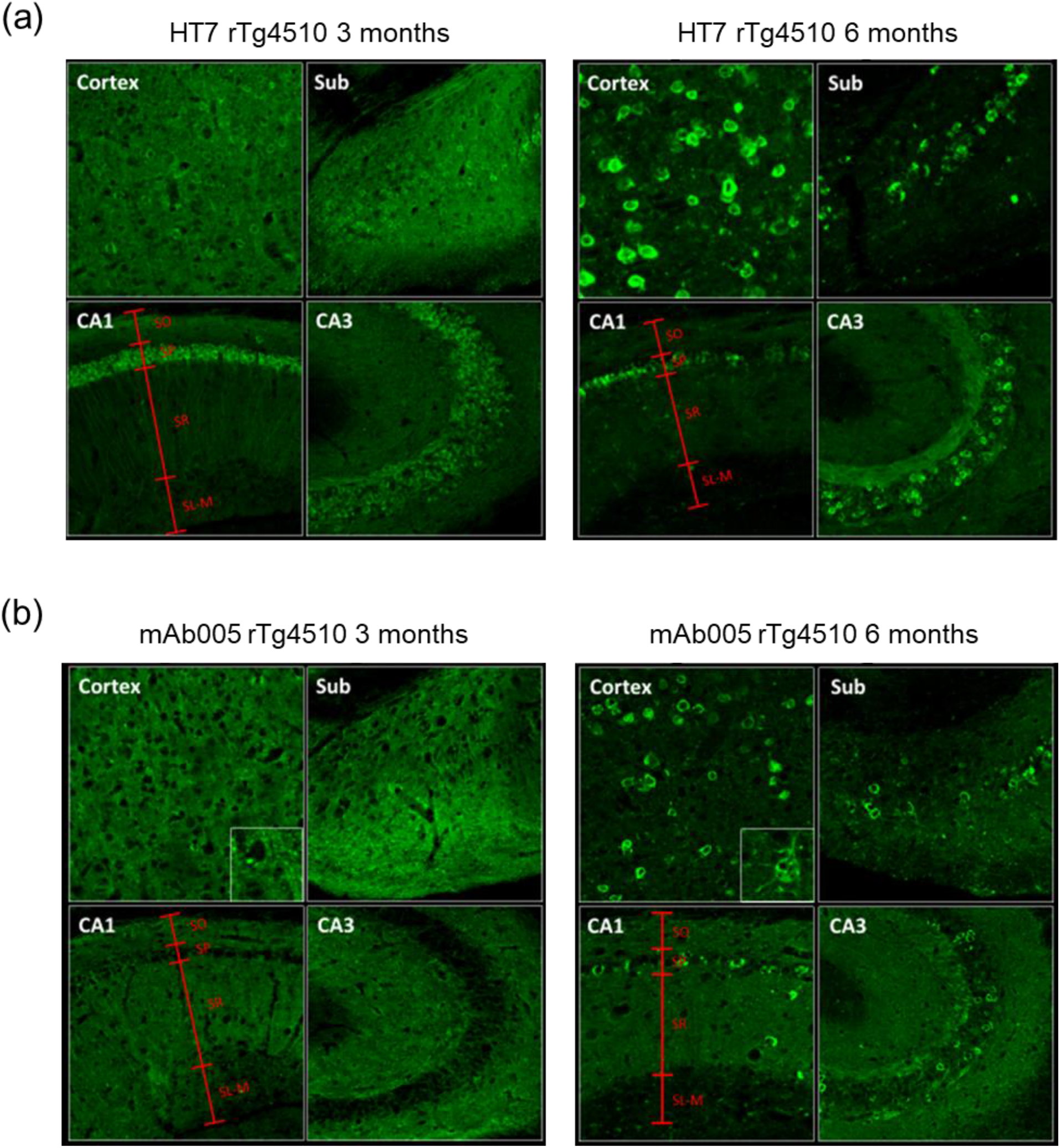
(a) The spatial distribution of overexpressed human tau (detected by HT7) in rTg4510 mice, at 3 and 6 months old. The neocortex, subiculum (Sub), hippocampal CA1 and CA3 regions are shown. In CA1 at 3 months, tau is highly expressed in neuronal somas in stratum pyramidale (SP) and relatively weak in the neuropil layers of stratum oriens (SO), stratum radiatum (SR), and stratum lacunosum moleculare (SLM). (b) mAb005 reactivity is very low in neuronal somas at 3 months, especially in the SP layer of CA1 and CA3, but relatively strong in the neuropil layers. At 6 months, mAb005 reactivity is strong in neuropils and the somas of tangle-bearing neurons.

In postmortem tissues, mAb005 was surprisingly unreactive against the abundant tau aggregates that accumulate by the end stage of AD, which include neurofibrillary tangles, ghost tangles, and neuropil threads. In the prefrontal cortex of AD subjects, mAb005 immunoreactivity was prevalent in Braak stage III/IV cases but largely absent in stage V/VI cases (**Fig. 4**). In the hippocampal CA1 region, mAb005 immunoreactivity was detectable in Braak stages III-V but largely vanished by stage VI (**Fig. S3**). By contrast, AT8, mAb004, mAb020, and mAb037 showed very strong staining during Braak stages V and VI in both areas. This early-detection feature distinguishes mAb005 from other known immuno-markers of pathological tau species. Therefore, we refer to the conformers recognized by mAb005 as early-stage oligomers of tau (esoTau) with respect to AD progression, which appear before widespread tangle deposition but disappear with tangle maturation.

**Figure 4.**
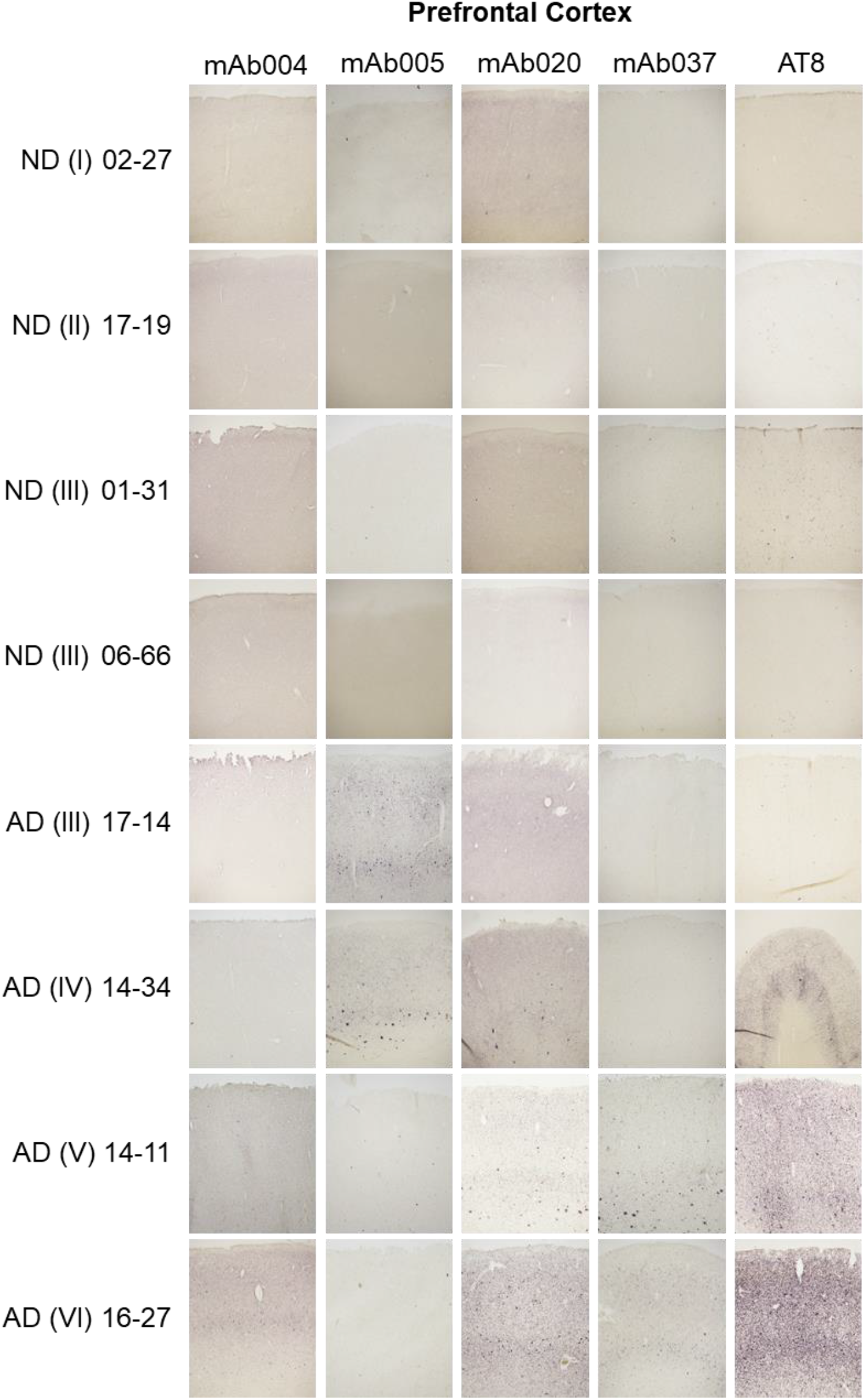
Immunohistochemistry of tau monoclonal antibodies (mAb series) and AT8 in the prefrontal cortex of AD subject (Braak stages III to VI) and non-demented (ND) subjects (Braak stages I to III). mAb005 immunoreactivity is prevalent in Braak stage III/IV AD subjects but largely missing in stage V/VI AD subjects. AT8 immunoreactivity increases with advanced Braak stages.

### mAb005 immunoreactivity in 3R and 4R tauopathies

Due to alternative splicing, tau protein may carry three or four microtubule repeats (3R or 4R) (*37*). AD is a mixed 3R and 4R tauopathy, and the tangle maturation process exhibits 4R-to-3R transition (*50*). Therefore, we investigated if mAb005 preferentially recognized 4R over 3R aggregates. We observed that mAb005 could recognize tau aggregates in 3R tauopathies (Pick’s disease, PiD) as well as 4R tauopathies (corticobasal degeneration, CBD, and progressive supranuclear palsy, PSP) (**Fig. S4**). In PSP subjects, mAb005 immunoreactivity was found within the tufted astrocytes of putamen, the oligodendrocytes of globus pallidus, and the neurons of nucleus basalis of Meynert **(Fig. 5**). Our results suggest that the mAb005 epitope is shared by 3R and 4R aggregates. The disappearance of mAb005 reactivity in Braak stage VI subjects may be related to conformation changes during tangle maturation rather than 4R-to-3R replacement. In sum, the misfolding event detected by mAb005 represents a common pathway associated with 3R, 4R, and mixed 3R/4R tauopathies, which may occur within synapses, neurons, astrocytes, or oligodendrocytes, depending on disease types and stages.

**Figure 5.**
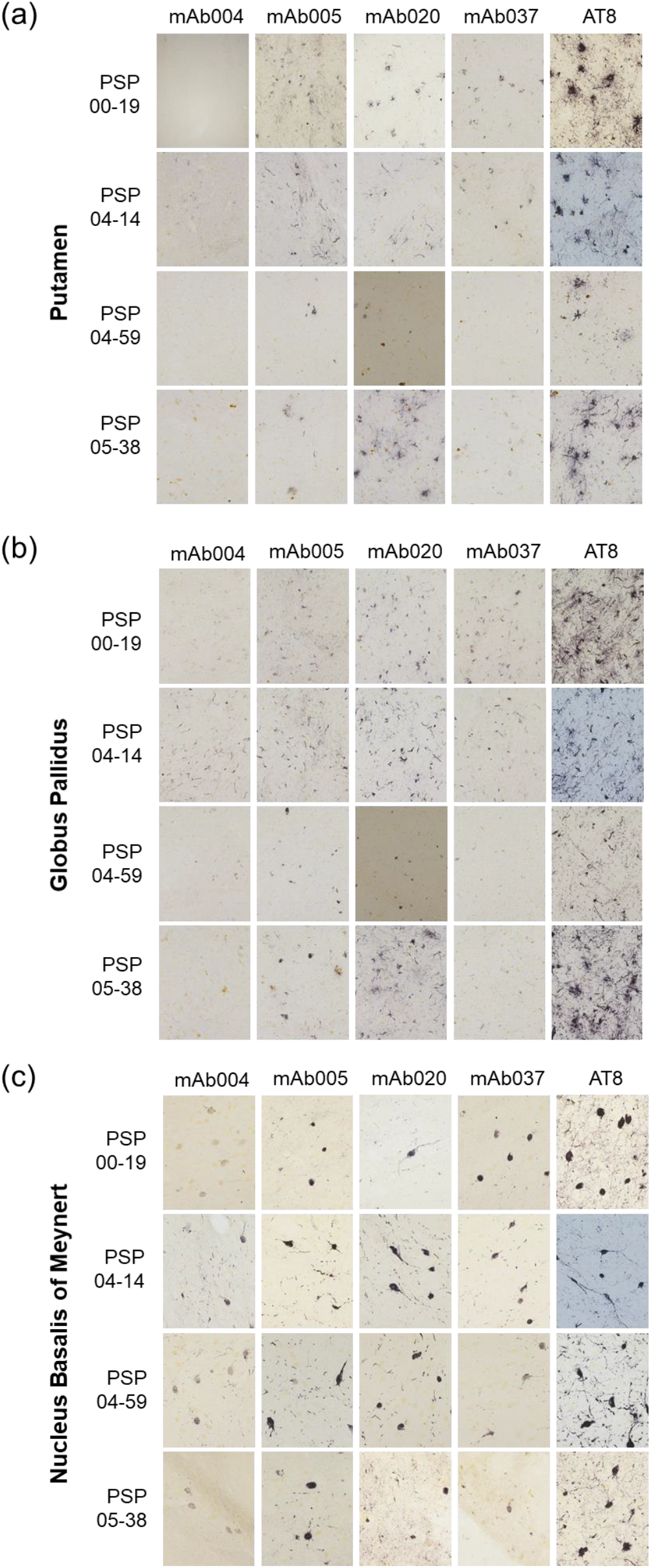
Immunohistochemistry of tau monoclonal antibodies (mAb series) and AT8 against different brain regions of PSP subjects: (a) putamen; (b) globus pallidus; (c) nucleus basalis of Meynert.

### Therapeutic effects of mAb005

In cellular models of uptake transmission, in which rTg4510 brain lysates were added to the culture media of cells expressing green fluorescent protein (GFP)-fused tau, mAb005 was effective in blocking intracellular GFP-tau aggregation with the EC_50_ of 72 nM (**Fig. 6a**). Therefore, blocking the uptake of toxic tau seeds is a plausible MOA for mAb005 immunotherapy. In rTg4510 mice receiving twelve weeks of high-dose mAb005 injections (50 mg/kg/week), we observed partial rescue of neuronal loss in the hippocampus dentate gyrus and CA1 region. At a lower dose of 10 mg/kg, neuronal loss was still reduced in the dentate gyrus (**Fig. 6b**). Total tau levels in brain lysates showed significant increases with both doses, which may be attributed to the enhanced survival of tau-expressing neurons. The levels of pTau (detected by AT8), synaptophysin, and esoTau (detected by mAb005) also increased in the high-dose group (**Fig. 6c**). mAb005 did not appear to promote the overall clearance of misfolded tau aggregates but may instead alleviate tau toxicity toward synapses and neurons.

**Figure 6.**
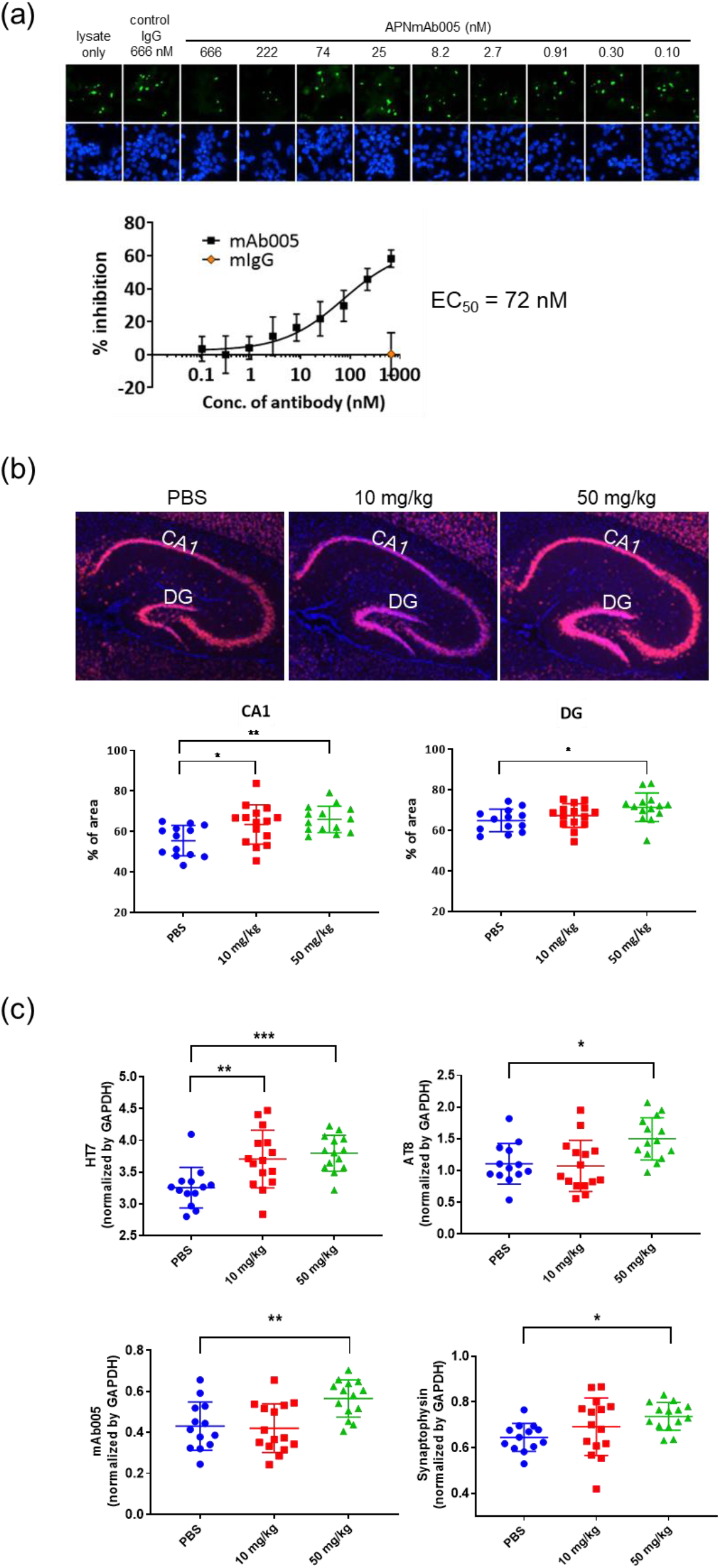
(a) rTg4510 brain lysate induces tau-GFP aggregation (green channel) in HEK cells (blue channel, DAPI staining), which is partially blocked by pre-incubation of mAb005 with brain lysates. The inhibition curve is shown with error bars representing standard deviations (n=3). (b) mAb005 injection partially rescues neuronal loss in the CA1 and dentate gyrus of rTg4510 mice, as quantified by % area of NeuN-positive staining in stratum pyramidale. (c) Quantification of total human tau (HT7), pTau (AT8), esoTau (mAb005), and synaptophysin levels in the brain lysates of rTg4510 mice injected with mAb005 antibody (n=13-15 per group; one-way ANOVA with Dunnett’s test: *p<0.05; **p<0.01, ***p<0.005).

## DISCUSSION

### Targeting synaptic tau oligomers in AD

The pathogenesis mechanisms of AD remain incompletely understood, but synaptic pathology may play important roles (*51-53*). Cognitive decline in AD is strongly correlated with synaptic loss, more so than neuronal loss, plaque counts, or tangle counts (*54, 55*). Multiple studies have implicated diffusible misfolded Aβ and tau species as key agents of synaptotoxicity (*53, 56, 57*). Moreover, the toxicity of Aβ on synapses may depend on tau as a downstream mediator (*58, 59*). We observed that Aβ induces pathological hyperphosphorylation patterns on tau proteins at synaptic sites (*60*), which may contribute to the accumulation of pTau oligomers inside the synapses of AD subjects (*34, 35*). Synaptic pTau proteins are also found in amyotrophic lateral sclerosis (*61*) and canine cognitive impairment (*62*), although their oligomerization states remain to be confirmed. According to the prion-like propagation hypothesis, synaptic pTau oligomers may be a crucial transmissive agent (*28-32*). Tau may be released extracellularly via synaptic vesicles (*63, 64*) or exchanged via exosomes, ectosomes, or nanotubes (*37, 65*). Therefore, we considered synaptic tau oligomers to be critically toxic and potentially transmissive, which makes them a promising target for tau immunotherapy.

Recent cryo-electron microscopy studies have identified over a dozen distinct conformations in tau fibrils associated with different tauopathies (*18, 66*). Although the conformations of tau oligomers remain to be resolved, cross-seeding studies have demonstrated the existence of different oligomer strains (*21, 67*), suggestive of different conformations. However, there is little understanding of how different oligomer strains are associated with different tauopathies at various respective stages. Conformation-specific antibodies remain indispensable tools for understanding the roles of oligomers in the pathogenesis of tauopathies (*68-72*).

Previously, we tested Alz-50 and MC-1 conformer antibodies side-by-side against AD-patient synaptosomes and observed positive staining for Alz-50 (*35*) but negative results for MC-1 (unpublished data). However, Alz-50 is less suitable for immunotherapy due to its IgM structure (*73*). Hence, we sought the development of IgG antibodies against synaptic tau oligomers. Our novel strategy of immunizing mice with AD synaptosome biomimetics (vesicles containing tau oligomers) led to a novel class of conformation-specific IgGs reactive against tau oligomers in AD-affected synapses (**Fig. 1e**). Among these, mAb005 was especially interesting due to two unique properties: (1) specificity against early-stage aggregates in AD subjects that largely disappeared by Braak stage VI (**Fig. 4 and S3**); (2) specificity for early diffuse misfolded species originating in distal neurites in mouse tauopathy models (**Fig. 3**). In rats with amyloid precursor protein knock-in, mAb005 immunoreactivity precedes MC-1 immunoreactivity by ∼10 months (*74*). While MC-1 antibody is thought to recognize misfolding events occurring between pretangle formation and tangle deposition (*75*), mAb005 appears to recognize even earlier misfolding/oligomerization events that generate what we termed esoTau.

### Comparisons against other conformational antibodies

In postmortem tissues of AD subjects, the immunoreactivity of Alz-50, MC-1, and TOC1 (an IgM against tau oligomers (*72*)) continue to accumulate from Braak stage IV to VI (*76, 77*). Likewise, pTau species detected by AT8 (pS202/pS205), PHF-1 (pS396/pS404), and CP13 (pT231) antibodies also peak at Braak stage VI (*76*). We do not yet understand why esoTau would disappear by late-stage AD, because no other pathological tau markers have shown comparable temporal patterns. One possibility is that esoTau eventually becomes incorporated into fibrillar structures and converts into other conformers. Another possibility is that esoTau may still be removed by chaperones or proteolytic systems, representing a reversible form of early-stage pathology.

In rTg4510 mice, MC-1 and TOC1 show perinuclear staining at 2.5 months (*78*), while TOMA (an IgG against tau oligomers) shows perinuclear staining at 4 months (*71*). The somatic clustering of tau aggregation is rightfully expected because tau expression is concentrated in neuronal somas in this transgenic model. In stark contrast, mAb005-reactive aggregates appear to be driven by a distinct misfolding mechanism occurring in distal neurites or around synapses, where fewer tau proteins are available. It is plausible that different cellular compartments may have different factors that drive tau misfolding, or that different misfolding species may be transported to different subcellular compartments.

It has been shown that tau aggregates from AD, PiD, CBD, and PSP subjects represent different transmissive strains (*79, 80*) and exhibit distinct fibril conformations (*18, 81-83*). Thus, we were surprised to find that mAb005 recognized tau aggregates in 3R tauopathies (PiD), 4R tauopathies (CBD and PSP) (**Fig. 5**), as well as mixed 3R/4R tauopathies (AD). It implies that the mAb005 epitope may situate outside the β-rich core of fibrillar tau, for which cryo-electron microscopic structures are available. EsoTau conformers do not appear to be “off-pathway” intermediates that temporarily occur in AD-affected neurons. Instead, esoTau conformers may represent oligomeric seeds found in diverse tauopathies and cellular compartments (neuronal somas, neuropils, synapses, astrocytes, and oligodendrocytes). The pervasive nature of esoTau makes it an attractive therapeutic target.

The therapeutic effects of conformation-specific antibodies in animal tauopathy models have been demonstrated, but the MOA may depend on the nature of the antibody and animal pathophysiology. For instance, MC-1 treatment effectively promoted the clearance of soluble and insoluble tau species in JNPL3 mice (P301S tau) (*84*), which may reflect its affinity for both smaller aggregates and larger fibrils (*85*). On the other hand, TOMA only reduced oligomer levels but not tangle or tau monomer levels in JNPL3 mice (*71*). In rTG4510 mice, which have more aggressive pathologies than JNPL3, TOMA did not affect tangle or tau monomer levels. It failed to rescue neuronal death in the hippocampus and behavioral deficits, but only managed to promoted the clearance of tau oligomers into the cerebrospinal fluid and blood serum (*86*). In comparison, APNmAb005 treatment promoted neuronal survival in the hippocampus of rTg4510 mice, accompanied by increases in total tau, pTau, synaptophysin, and esoTau levels (**Fig. 6**). The MOA of mAb005 may not be general tau clearance but the neutralization of certain misfolded species that are toxic against synapses and neurons. Whether such neutralization occurs intracellularly or extracellularly remains unclear.

## CONCLUSION

We successfully generated a group of conformational IgG antibodies that recognized synaptic tau oligomers from AD subjects. Among these, APNmAb005 showed remarkable specificity against misfolded conformers and was selected for detailed characterization. mAb005 recognized tau aggregates in early-stage AD but not the abundant inclusions at the end stage. In rTg4510 mice, mAb005 reactivity first developed in distal neurites but not somas. These results suggest that different types of tau oligomers may accumulate with distinct spatiotemporal patterns. In human brains, mAb005 immunoreactivity was found in both nerve and glial cells, as well as in 3R, 4R, and mixed tauopathies. Its target appears to be early-stage oligomeric seeds shared by many tauopathies, which we termed esoTau.

In cell cultures, mAb005 blocked the cellular uptake of toxic tau species contained in rTg4510 brain lysates. *In vivo*, mAb005 injection into rTg4510 mice promoted neuronal survival and preserved synaptic integrity. It appears to neutralize the toxic effects of early-misfolding tau species (esoTau) without apparent clearance of tau aggregates. Targeting early-stage tau oligomers around synaptic compartments represents a novel strategy for tau immunotherapy. A humanized version of mAb005 has undergone scaled-up production and entered phase 1 clinical trial.

## MATERIALS AND METHODS

### Reagents

The following antibodies were purchased commercially: HT7 (Thermo MN1000, AB_2314654), rabbit anti-tau (DAKO A0024, AB_10013724), AT8 (pS202/pT205, Thermo MN1020, AB_223647), synaptophysin clone SY38 (Abcam ab8049, AB_2198854; GeneTex GTX72051, AB_373523) and GAPDH (Proteintech 10494-1-AP, AB_2263076). Secondary antibodies were purchased from LI-COR (926-32210, 926-68071, and 926-68078) and Jackson ImmunoResearch (115-606-146). Dithiothreitol (DTT), sodium dodecyl sulfate (SDS), and HEPES were purchased from Sigma.

### Generation of tau monoclonal antibodies

All anti-tau monoclonal antibodies were generated by the hybridoma approach against recombinant tau aggregates (human full-length tau) encapsulated in the ACM Polymersomes (ACM Biolabs, Singapore) and purified by size exclusion chromatography (SEC) to remove the non-encapsulated aggregates. The inclusion of the tau aggregates was confirmed by the specific binding of the fluorescent dye PBB3 (*87*). Standard hybridoma approach was conducted to immunize the wild-type BALB/c and SJL mice with 3 rounds of 5 or 10 µg of antigens. ELISA assays were performed using recombinant tau aggregates to check the degree of immunization and subsequent steps of clone selection and expansion. Approximate 50 antibodies were further purified and characterized by immunoblotting for tau binding. Eleven antibodies were selected based on their tau binding patterns correlating well to pathological tau species characterized in the literature.

### Postmortem tissues and human subjects

All human brain tissues of non-demented (ND), AD, PSP, PiD, and CBD subjects were obtained from Banner Sun Health Research Institute (Sun City, AZ, USA) (*88*). Brain tissues for immunoblotting and dot blotting were processed in house (at APRINOI) as described in designated sections. Pathological scoring of tau immunohistochemical binding was conducted at Banner Sun Health Research Institute accordingly. All donor tissues were obtained in accord with local and national IRB regulations.

### Brain total lysate

Approximately 200 mg of human brain tissue was dissected out by a microtome blade on the dry ice block, and disrupted in ice-cold Tris-based lysis buffer (50 mM Tris, 2 mM EGTA and EDTA, 274 mM NaCl, 5 mM KCl, protease and phosphatase inhibitors, pH 8.0) using lysing matrix D beads (MP Biomedicals, 6913-500) by the FastPrep homogenizer (MP Biomedicals). The homogenate was spun for 15 min at 13,000 xg at 4°C, and the supernatant was collected as brain total lysate. The protein concentration was determined by Bradford assay (BioRad).

### Sarkosyl extraction

Brain total lysate was incubated in Tris-based lysis buffer containing 1% sarkosyl (diluted from 10% sarkosyl stock buffer, Bioworld, 41930024-4) at room temperature (RT) for 1 hr, and spun at 150,000 xg for 20 min at 4 °C. The supernatant was collected as the sarkosyl-soluble fraction (S1). The remaining pellet was resuspended in phosphate buffered saline (PBS) and spun at 150,000 xg for 20 mins at 4°C. The supernatant was removed and the pellet was resuspended in PBS as the sarkosyl-insoluble fraction (P2).

### Synaptosome preparation

Brain tissues were dissected into small pieces (∼300 mg) by microtome blade, and then disrupted by Teflon-glass homogenizer in ice-cold buffer A (25 mM HEPES, 120 mM NaCl, 5 mM KCl, 1 mM MgCl_2_, 2 mM CaCl_2_, protease and phosphatase inhibitors, 2 mM DTT, pH 7.5). The homogenate was passed through an 80-µm filter (Millipore, NY8002500) and a 5-µm filter (Pall, 4662). The filtrate was spun at 1,000 xg for 10 mins at 4 °C. The pellet was labeled as P1 (synaptosomes). The supernatant was further spun at 100,000 xg for 30 mins at 4 °C to remove organelles and microsomes, and the supernatant was collected as the cytosol fraction. An aliquot of P1 was incubated with 0.5% Triton-X 100 (Thermo, 28314) in PBS at 4 °C and rotated for 20 min. After centrifugation at 15,000 xg for 15 min at 4 °C, the supernatant was collected as synaptosome extract for dot-blotting. Another aliquot of P1 was resuspended in buffer A and spun at 1,000 xg for 10 mins at 4 °C to yield the P1’ pellet. This was resuspended in buffer B (50 mM Tris, 1.5 % SDS, 2 mM DTT, pH 7.5) and boiled for 5 min, followed by centrifugation at 15,000 xg for 15 min. The supernatant was collected as synaptosome extract for immunoblotting.

### Immunoblotting and dot blotting

Brain total lysate was supplemented with 50 mM DTT and LDS sample buffer (Thermo, B0007) and boiled for 5 min. It was resolved by 4-12 % gradient Bolt Bis-Tris Plus gels (Thermo, NW0415BOX), and transferred onto a PVDF membrane. The membrane was incubated in blocking buffer (LI-COR, 927-50000) for 30 min. Primary antibodies were incubated for 1 hr at RT, followed by three washes in PBS. Secondary antibodies were incubated for 1 hr at RT, followed by three washes in PBS. Imaging and quantification of signals were performed by LI-COR Odyssey infrared imaging system.

For dot blots, non-denatured brain total lysate (0.4 µg protein), sarkosyl-soluble and insoluble fractions (10 ng of tau), and synaptosomal and cytosolic extracts (0.4 µg protein) were adjusted to 2 µL in volume and spotted onto a nitrocellulose membrane (Thermo, LC2001). The membrane was blocked for 30 min, incubated with primary antibodies for 1 hr, and incubated with secondary antibodies for 1 hr.

### Size exclusion chromatography

Brain total lysate (1 mg protein) was spun at 15,000 xg for 10 mins to remove insoluble materials. The supernatant was collected and loaded into Superose 6 Increase column (Cytiva, 29-0915-96) and eluted in 30 mM Tris buffer (pH 7.4). Gel filtration standards (Bio-Rad, 1511901) were used to determine the molecular weight of proteins in each fraction.

### Sucrose gradient centrifugation

A discontinuous gradient containing 10, 20, 30, 40, and 50% sucrose solutions (1.5 ml each) were layered into a centrifuge tube (Beckman, 331372). Brain total lysate (6 mg of protein, 3 ml) was carefully added to the top of the sucrose gradient, and spun at 250,000 xg in a SW41 rotor (Beckman) for 4 hr at 4 °C. Each layer (top, 10, 20, 30, 40, and 50% sucrose fractions) was carefully collected from top to bottom for further analysis.

### Immunohistochemistry for human brains

To determine the tissue constituents stained in normal brain tissues from clinically diagnosed ND, AD, and PSP elderly subjects (**SI Table S1**), two to four sections were stained from each brain. AD subjects included two at Braak stage III-IV and two at Braak stage V-VI, while ND subjects included two at Braak stage III and one each at Braak stages I and II. Subjects with PSP are not assigned a Braak stage as the tau pathology is not of the AD type. PSP cases were chosen to represent typical PSP with few or no neuritic (amyloid) plaques. Brain regions stained for both AD and PSP cases were the frontal lobe: including superior frontal gyrus, middle frontal gyrus, inferior frontal gyrus, and anterior cingulate gyrus. AD cases were additionally stained for the temporal lobe: including amygdala, entorhinal and transentorhinal areas, hippocampus, dentate gyrus, and parahippocampal gyrus with entorhinal and transentorhinal areas, fusiform gyrus, inferior, middle, and superior temporal gyrus. PSP cases were additionally stained for the basal ganglia: including putamen, globus pallidus, and nucleus basalis of Meynert. ND control subjects had sections stained to correspond with those stained in both AD and PSP groups.

Formalin-fixed, free-floating sections (40 μm) were first treated with 1% H_2_O_2_ (Thermo). After 30 min, the tissue was incubated with APNmAb005 antibody (Lot 170925005) (1:10,000 for AD sections; 1:1,000 for PSP sections) and AT8 (1:1,000) for 24 hours with agitation. The tissue was washed by PBS-TX (0.3% Triton X-100) and then incubated with 1:1000 dilution of biotinylated anti-mouse antibody (Vector Labs) with agitation. After 2 hours, the tissue was incubated with avidin-biotin peroxidase complex (Vector Labs) for 30 min with agitation. The tissue was then incubated with DAB and nickel ammonium sulfate for 25 min with agitation and moved into Tris-buffer (pH 7.5) for 25 mins. After cell, fiber, and tangle signals become clearly visible, the tissue was stained with Neutral Red and then mounted with Permount mounting solution (Thermo).

### Therapeutic effects in rTg4510 mice

rTg4510 mice were licensed from Mayo Clinics and housed in National Laboratory Animal Center (Taipei, Taiwan) at 24°C with a 12 -hr light/dark cycle. Food and tap water were provided ad libitum. Around 3 months of age, male mice were transferred to Taiwan Mouse Clinics (Taipei, Taiwan) for 12-week dosing study (IACUC No. 13-5-763). All animals were maintained at 21 ± 2°C, 50 ± 5% humidity, with a 12-hr light/dark cycle. Food and tap water were provided ad libitum.

A 7-day acclimation period was implemented prior to drug dosing. On the dosing day, all animals were weighed before drug administration and assigned into 3 groups by random sampling method. Mice received twice-daily intraperitoneal dosing (i.p.) of 5 µl/g bodyweight of APNmAb005 (10 or 50 mg/kg bodyweight) or vehicle (PBS) with insulin syringe. Adequate measures were utilized to minimize potential pain or discomfort experienced by the mice used in this study. All the experiments involving animals adhered to relevant ethical guidelines and the ARRIVE guidelines.

### Immunohistochemistry for mouse brains

The hemibrains were sliced by Cryomicrotome (Leica) and examined at the pathology core in the Institute of Biomedical Sciences, Academia Sinica (Taipei, Taiwan). Each hemibrain was mounted in cryo-embedding media (OCT) on the base mold on dry ice until the tissue was completely frozen and stored at -80°C. All frozen tissue blocks were sectioned within a week. The frozen tissue block was sliced into sagittal 30-µm cyro sections by a cryotome cryostat at -20°C. The sequential sections contained horizontal parts of the cerebrum, hippocampus, and cerebellum between +0.24 to +2.64 mm laterally from the bregma. Each brain slice was kept in a microcentrifuge tube with 1 ml PBS at 4°C.

The slices were washed three times in 1.5 ml PBS for 10 min each, and incubated in PBS with 0.3% Triton-X100 for 1 hr at RT. The slices were blocked in blocking buffer (10% normal goat serum and 5% BSA in PBS), incubated with goat anti-mouse IgG Fab fragment (Jackson ImmunoResearch, 115-007-003, 20 ug/ml in PBS) for 1 hour at RT, and washed three times to thoroughly remove the Fab. This was followed by incubation with primary antibodies overnight at 4 °C and three washes. Incubation with secondary antibodies for 1 hour at RT was followed by three washes. The slices were transferred to new wells with DAPI staining for 10 minutes at RT and washed again. Afterwards, the slices were transferred onto microslides (Muto Pure Chemicals, 1-6724-01) and sealed with mounting medium (Thermo, P36930). The coverslip (Dogger, AP-0101050) was applied and sealed with nail polish.

### Tau seeding inhibition assay

Brains were obtained from 6-month-old rTg4510 mice and the cerebellum was removed by microtome blade. The remaining brain tissue was disrupted in Tris-based lysis buffer (50 mM Tris-base, 2 mM EGTA and EDTA, 274 mM NaCl, 5 mM KCl, protease and phosphatase inhibitors, pH 8.0) using lysing matrix D beads. The homogenate was spun for 15 mins at 13,000 xg at 4 °C. The supernatant was collected as brain total lysate and stored at -80 °C. The total lysate was exchanged into PBS buffer using centrifugal filter devices (Merck, UFC201024) according to manufacturer protocols.

rTg4510 brain lysate in PBS was incubated with serially diluted APNmAb005 or control mouse IgG for 2 hr at RT with rotation. The antibody-lysate mixture was added to HEK cells stably expressing TauLM-GFP fusion protein (full length 4R tau with P301L and V337M mutations). After 48 hours at 37°C, the cells were fixed by 4% paraformaldehyde (Electron Microscopy Sciences, 15710) and stained with 10 nM of DAPI. Images were acquired by ImageXpress Micro Confocal (Molecular Devices) and further analyzed to quantify the number of cells (GFP and DAPI double positive) and tau spots.

### Preparation of tau oligomers

Full-length tau was expressed in *E. coli* (One Shot BL21 Star DE3, Thermo) using the pRK172-htau40 plasmid. Bacteria were lysed in 20 mM Tris buffer (pH 9.0) by sonication. The lysate was boiled for 10 min to precipitate heat-unstable proteins (tau remained soluble) and sonicated again. The lysate was cooled to 4 °C and cleared by spinning at 15,000 xg for 20 min. The supernatant was subjected to anion exchange chromatography (Cytiva HiTrap Q HP column) with gradient elution (0-250 mM NaCl). Tau-containing fractions were collected and subjected to SEC (Cytiva HiLoad 16/600 superdex 200 pg column) and eluted in 20 mM Tris buffer (pH 7.4), yielding highly purified tau monomers. Tau protein aggregation was induced with 0.3 mg/mL heparin in 20 mM MOPS buffer (pH 7.0) at 37 °C. The fibrillar and oligomeric fractions were isolated by SEC.

### Statistical Analysis

EC_50_ and ANOVA were calculated using GraphPad Prism 8 software (San Diego, CA, USA).

## Acknowledgement

The htau40 plasmid was generously provided by Michele Goedert, Medical Research Council Laboratory. We thank Ying-Ju Wu and Yung-Hsiang Hsu for preparing tau oligomer samples. We thank Taiwan Mouse Clinics, National Laboratory Center and Charles River Laboratories for their kind assistance in dosing, breeding, and rTg4510 mouse immunohistochemistry. We are grateful to the Banner Sun Health Research Institute Brain and Body Donation Program of Sun City, Arizona for the provision of human biological materials.

## Author contributions

Conceptualization: HCT, CYT, and MKJ. Methodology and Investigation: HCT, HTM, SSH, MFW, CLW, YTL, CLL, RM, AJI, GES, TGB, MN, BN, and CYT. Visualization: HCT, TGB, and CYT. Supervision: CYT and MKJ. Writing – original draft: HCT and CYT. Writing – review and editing: TGB, BN, and MKJ.

## Data and Materials Availability

All data are available from the authors on reasonable request.

## List of Supplementary Materials

Table S1

Figure S1 to S4.

## Supplementary Materials

**Table S1.** Human subjects for neuropathological examination.

| Diagnosis | Case # | Age | Sex | Braak Stage | Plaque Density |
| --- | --- | --- | --- | --- | --- |
| AD | 14-11 | 82 | M | V | frequent |
| AD | 14-34 | 97 | F | IV | frequent |
| AD | 16-27 | 94 | F | VI | frequent |
| AD | 17-14 | 95 | F | III | frequent |
| ND control | 01-31 | 81 | M | III | sparse |
| ND control | 02-27 | 86 | M | I | sparse |
| ND control | 06-66 | 78 | M | III | sparse |
| ND control | 17-19 | 53 | F | II | zero |
| PSP | 00-19 | 64 | M | n/a | zero |
| PSP | 04-14 | 83 | M | n/a | zero |
| PSP | 04-59 | 93 | M | n/a | moderate |
| PSP | 05-38 | 80 | M | n/a | zero |

**Figure S1.**
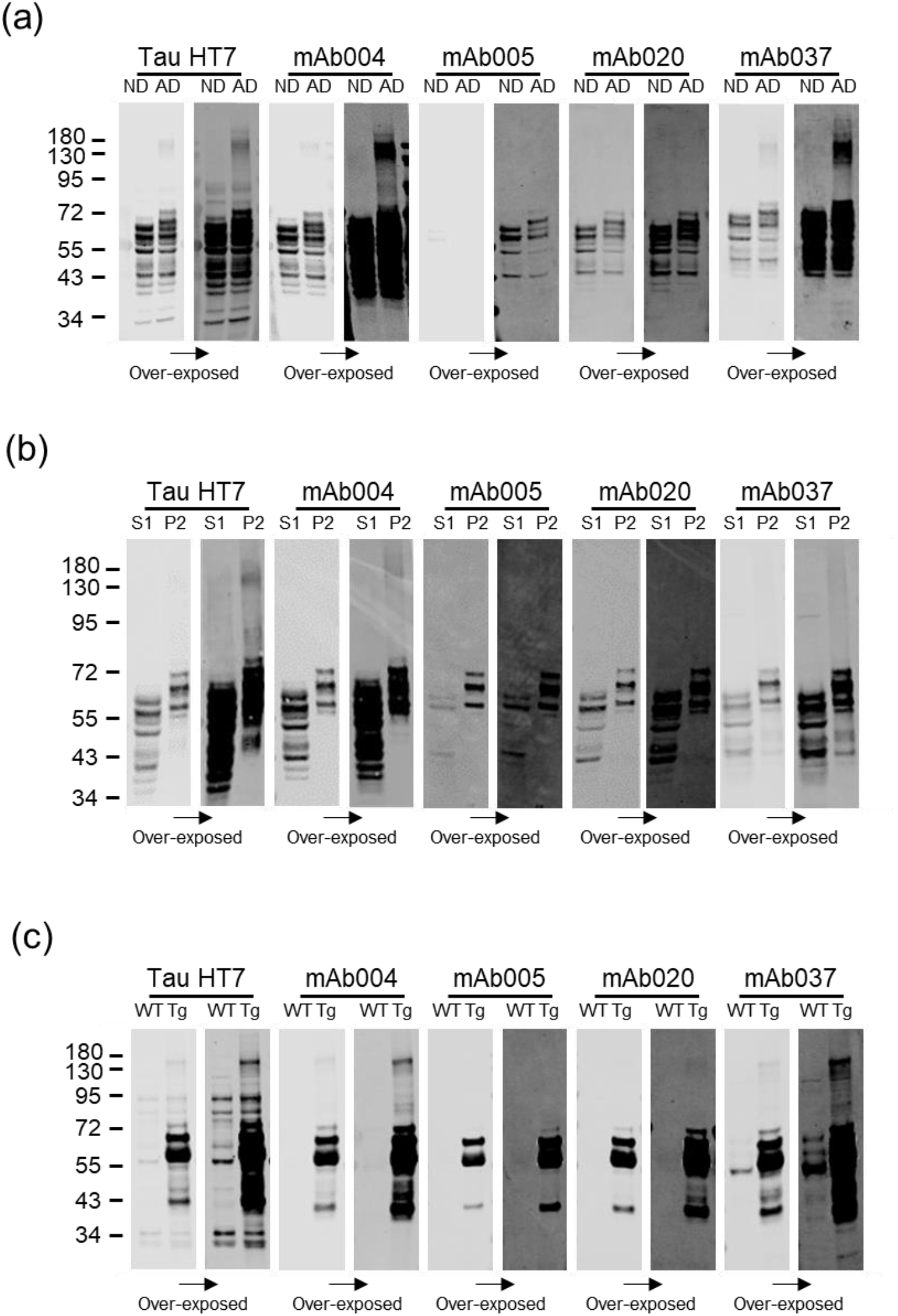
Immunoblotting of monoclonal tau antibodies (mAb series) against: (a) brain total lysates of non-demented (ND) and AD subjects; (b) sarkosyl-soluble (S1) and insoluble (P2) fractions of AD brain extracts; (c) brain total lysates of wild-type and rTg4510 mice.

**Figure S2.**
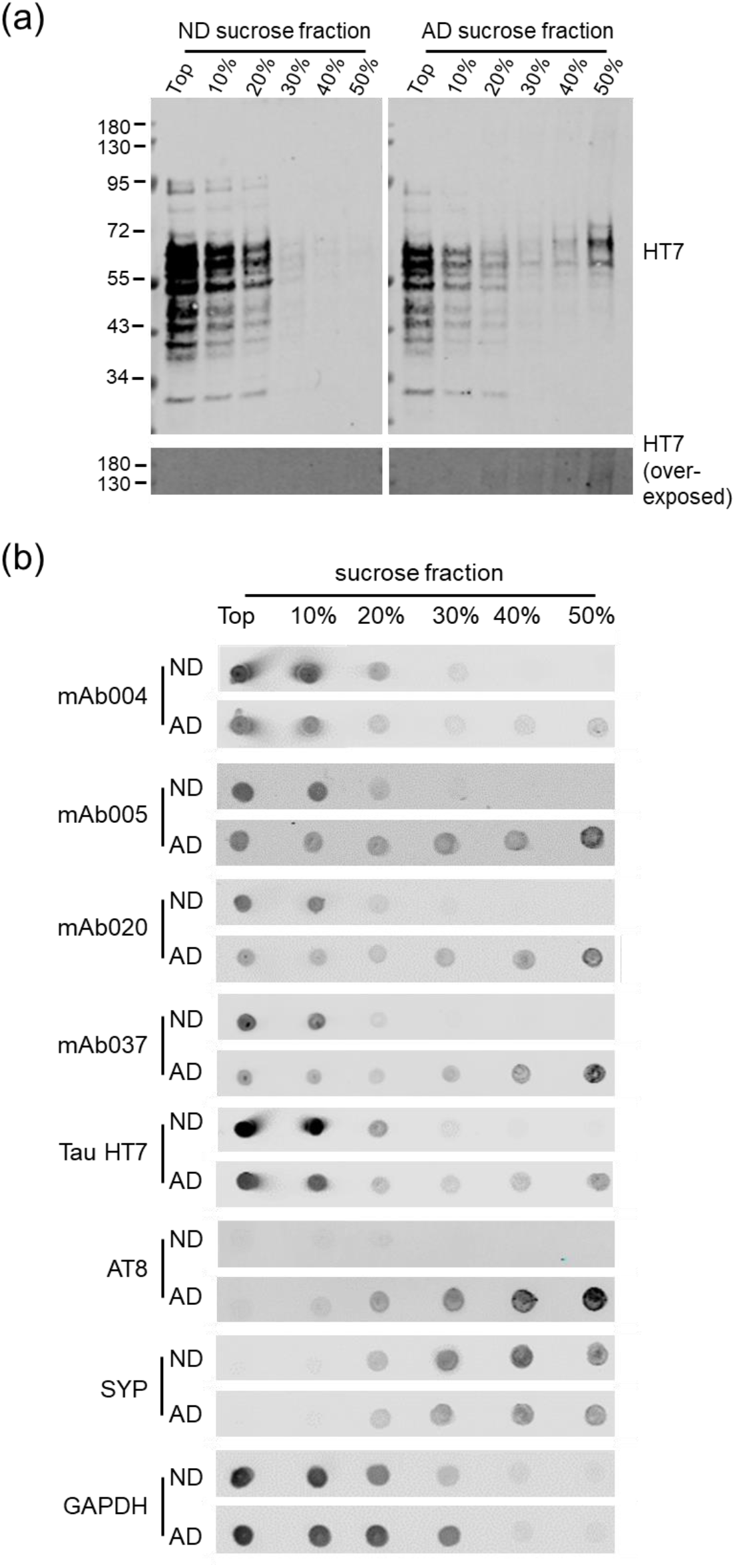
Discontinuous sucrose gradient centrifugation of brain extracts from non-demented (ND) and AD subjects, analyzed by: (a) SDS-PAGE and immunoblotting analysis against human tau (HT7); (b) Dot blotting using tau monoclonal antibodies (mAb series), total tau (HT7), pTau (AT8), synaptophysin (SYP), and GAPDH.

**Fig S3.**
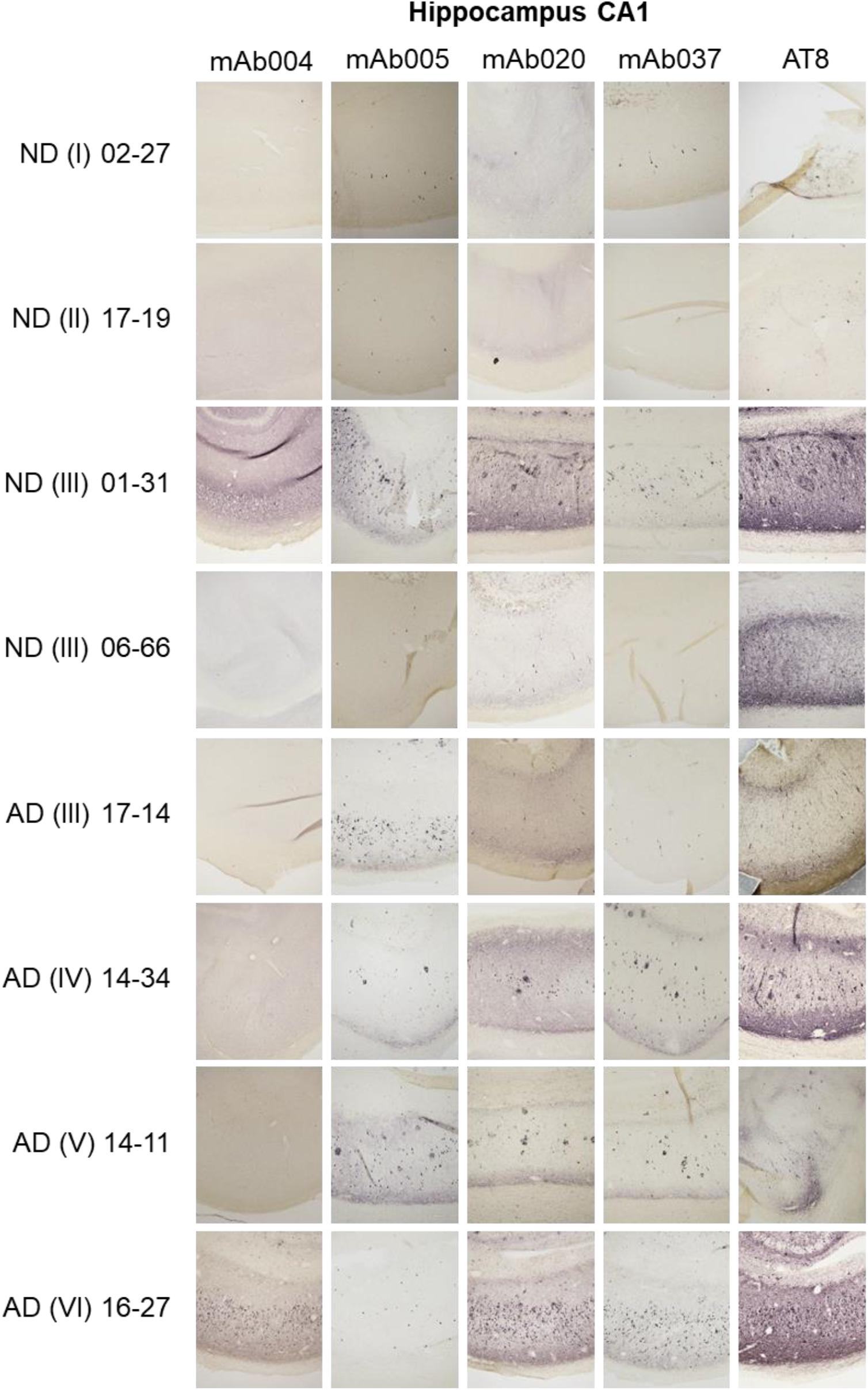
Characterization of tau monoclonal antibodies (mAb series) using hippocampal tissues of AD subjects and non-demented (ND) controls. mAb005 immunoreactivity is prevalent in Braak stage III-V AD subjects but very weak in Braak stage VI.

**Fig. S4.**
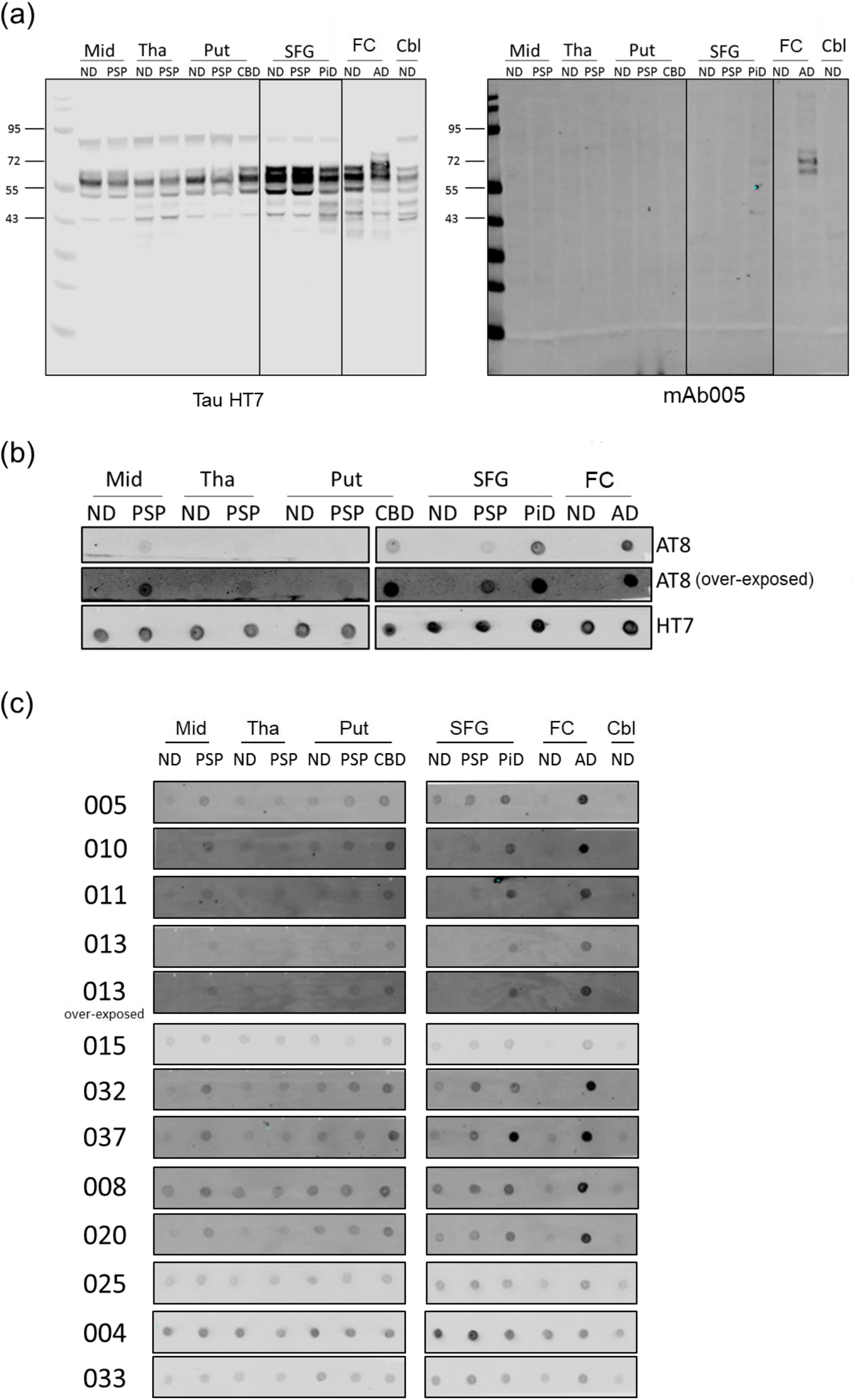
(a) Detection of tau species by human tau antibody (HT7) and mAb005 in human brain lysates. Regions analyzed: midbrain (Mid), thalamus (Tha), putamen (Put), superior frontal gyrus (SFG), frontal cortex (FC), and cerebellum (Cbl). The subjects included non-demented (ND), PSP, CBD, and AD cases. (b) Dot blot analysis against pTau (AT8). (c) Dot blot characterization for tau monoclonal antibodies.

## References

1. A. Serrano-Pozo, M. P. Frosch, E. Masliah, B. T. Hyman, Neuropathological alterations in Alzheimer disease. Cold Spring Harb. Perspect. Med. 1, a006189 (2011).

2. S. Spina, R. La Joie, C. Petersen, A. L. Nolan, D. Cuevas, C. Cosme, M. Hepker, J.-H. Hwang, Z. A. Miller, E. J. Huang, A. M. Karydas, H. Grant, A. L. Boxer, M. L. Gorno-Tempini, H. J. Rosen, J. H. Kramer, B. L. Miller, W. W. Seeley, G. D. Rabinovici, L. T. Grinberg, Comorbid neuropathological diagnoses in early versus late-onset Alzheimer’s disease. Brain 144, 2186–2198 (2021).

3. G. Niewiadomska, W. Niewiadomski, M. Steczkowska, A. Gasiorowska, Tau oligomers neurotoxicity. Life 11, 28 (2021).

4. C. Wells, S. Brennan, M. Keon, L. Ooi, The role of amyloid oligomers in neurodegenerative pathologies. Int. J. Biol. Macromol. 181, 582–604 (2021).

5. B. B. Holmes, J. L. Furman, T. E. Mahan, T. R. Yamasaki, H. Mirbaha, W. C. Eades, L. Belaygorod, N. J. Cairns, D. M. Holtzman, M. I. Diamond, Proteopathic tau seeding predicts tauopathy in vivo. Proc. Natl. Acad. Sci. U. S. A. 111, E4376–4385 (2014).

6. B. G. Perez-Nievas, T. D. Stein, H. C. Tai, O. Dols-Icardo, T. C. Scotton, I. Barroeta-Espar, L. Fernandez-Carballo, E. L. de Munain, J. Perez, M. Marquie, A. Serrano-Pozo, M. P. Frosch, V. Lowe, J. E. Parisi, R. C. Petersen, M. D. Ikonomovic, O. L. Lopez, W. Klunk, B. T. Hyman, T. Gomez-Isla, Dissecting phenotypic traits linked to human resilience to Alzheimer’s pathology. Brain 136, 2510–2526 (2013).

7. T. Athar, K. Al Balushi, S. A. Khan, Recent advances on drug development and emerging therapeutic agents for Alzheimer’s disease. Mol. Biol. Rep. 48, 5629–5645 (2021).

8. C. M. Vander Zanden, E. Y. Chi, Passive Immunotherapies Targeting Amyloid Beta and Tau Oligomers in Alzheimer’s Disease. J. Pharm. Sci. 109, 68–73 (2020).

9. D. Jeremic, L. Jimenez-Diaz, J. D. Navarro-Lopez, Past, present and future of therapeutic strategies against amyloid-beta peptides in Alzheimer’s disease: a systematic review. Ageing Res. Rev. 72, 101496 (2021).

10. C. Ji, E. M. Sigurdsson, Current Status of Clinical Trials on Tau Immunotherapies. Drugs 81, 1135–1152 (2021).

11. K. Y. Liu, R. Howard, Can we learn lessons from the FDA’s approval of aducanumab? Nat. Rev. Neurol. 17, 715–722 (2021).

12. J. Sevigny, P. Chiao, T. Bussière, P. H. Weinreb, L. Williams, M. Maier, R. Dunstan, S. Salloway, T. Chen, Y. Ling, J. O’Gorman, F. Qian, M. Arastu, M. Li, S. Chollate, M. S. Brennan, O. Quintero-Monzon, R. H. Scannevin, H. M. Arnold, T. Engber, K. Rhodes, J. Ferrero, Y. Hang, A. Mikulskis, J. Grimm, C. Hock, R. M. Nitsch, A. Sandrock, The antibody aducanumab reduces Aβ plaques in Alzheimer’s disease. Nature 537, 50–56 (2016).

13. M. Tolar, S. Abushakra, J. A. Hey, A. Porsteinsson, M. Sabbagh, Aducanumab, gantenerumab, BAN2401, and ALZ-801-the first wave of amyloid-targeting drugs for Alzheimer’s disease with potential for near term approval. Alzheimers Res. Ther. 12, 95 (2020).

14. E. M. Mandelkow, E. Mandelkow, Biochemistry and cell biology of tau protein in neurofibrillary degeneration. Cold Spring Harb. Perspect. Med. 2, a006247 (2012).

15. T. E. Tracy, L. Gan, Tau-mediated synaptic and neuronal dysfunction in neurodegenerative disease. Curr. Opin. Neurobiol. 51, 134–138 (2018).

16. J. Avila, J. J. Lucas, M. Perez, F. Hernandez, Role of tau protein in both physiological and pathological conditions. Physiol. Rev. 84, 361–384 (2004).

17. S. Banerjee, A. Ghosh, Structurally Distinct Polymorphs of Tau Aggregates Revealed by Nanoscale Infrared Spectroscopy. J. Phys. Chem. Lett. 12, 11035–11041 (2021).

18. Y. Shi, W. Zhang, Y. Yang, A. G. Murzin, B. Falcon, A. Kotecha, M. van Beers, A. Tarutani, F. Kametani, H. J. Garringer, R. Vidal, G. I. Hallinan, T. Lashley, Y. Saito, S. Murayama, M. Yoshida, H. Tanaka, A. Kakita, T. Ikeuchi, A. C. Robinson, D. M. A. Mann, G. G. Kovacs, T. Revesz, B. Ghetti, M. Hasegawa, M. Goedert, S. H. W. Scheres, Structure-based classification of tauopathies. Nature 598, 359–363 (2021).

19. S. Dujardin, C. Commins, A. Lathuiliere, P. Beerepoot, A. R. Fernandes, T. V. Kamath, M. B. De Los Santos, N. Klickstein, D. L. Corjuc, B. T. Corjuc, P. M. Dooley, A. Viode, D. H. Oakley, B. D. Moore, K. Mullin, D. Jean-Gilles, R. Clark, K. Atchison, R. Moore, L. B. Chibnik, R. E. Tanzi, M. P. Frosch, A. Serrano-Pozo, F. Elwood, J. A. Steen, M. E. Kennedy, B. T. Hyman, Tau molecular diversity contributes to clinical heterogeneity in Alzheimer’s disease. Nat. Med. 26, 1256–1263 (2020).

20. A. Kraus, E. Saijo, M. A. Metrick, 2nd, K. Newell, C. J. Sigurdson, G. Zanusso, B. Ghetti, B. Caughey, Seeding selectivity and ultrasensitive detection of tau aggregate conformers of Alzheimer disease. Acta Neuropathol. 137, 585–598 (2019).

21. F. Lo Cascio, S. Garcia, M. Montalbano, N. Puangmalai, S. McAllen, A. Pace, A. Palumbo Piccionello, R. Kayed, Modulating disease-relevant tau oligomeric strains by small molecules. J. Biol. Chem. 295, 14807–14825 (2020).

22. H. Wesseling, W. Mair, M. Kumar, C. N. Schlaffner, S. Tang, P. Beerepoot, B. Fatou, A. J. Guise, L. Cheng, S. Takeda, J. Muntel, M. S. Rotunno, S. Dujardin, P. Davies, K. S. Kosik, B. L. Miller, S. Berretta, J. C. Hedreen, L. T. Grinberg, W. W. Seeley, B. T. Hyman, H. Steen, J. A. Steen, Tau PTM Profiles Identify Patient Heterogeneity and Stages of Alzheimer’s Disease. Cell 183, 1699–1713 e1613 (2020).

23. C. Alquezar, S. Arya, A. W. Kao, Tau Post-translational Modifications: Dynamic Transformers of Tau Function, Degradation, and Aggregation. Front. Neurol. 11, 595532 (2020).

24. S. W. Min, S. H. Cho, Y. Zhou, S. Schroeder, V. Haroutunian, W. W. Seeley, E. J. Huang, Y. Shen, E. Masliah, C. Mukherjee, D. Meyers, P. A. Cole, M. Ott, L. Gan, Acetylation of tau inhibits its degradation and contributes to tauopathy. Neuron 67, 953–966 (2010).

25. C. Smet-Nocca, M. Broncel, J.-M. Wieruszeski, C. Tokarski, X. Hanoulle, A. Leroy, I. Landrieu, C. Rolando, G. Lippens, C. P. Hackenberger, Identification of O-GlcNAc sites within peptides of the Tau protein and their impact on phosphorylation. Mol. Biosyst. 7, 1420–1429 (2011).

26. S. N. Thomas, K. E. Funk, Y. Wan, Z. Liao, P. Davies, J. Kuret, A. J. Yang, Dual modification of Alzheimer’s disease PHF-tau protein by lysine methylation and ubiquitylation: a mass spectrometry approach. Acta Neuropathol. 123, 105–117 (2012).

27. E. B. Dammer, A. K. Lee, D. M. Duong, M. Gearing, J. J. Lah, A. I. Levey, N. T. Seyfried, Quantitative phosphoproteomics of Alzheimer’s disease reveals cross-talk between kinases and small heat shock proteins. Proteomics 15, 508–519 (2015).

28. C. Peng, J. Q. Trojanowski, V. M. Lee, Protein transmission in neurodegenerative disease. Nat. Rev. Neurol. 16, 199–212 (2020).

29. C. A. Lasagna-Reeves, D. L. Castillo-Carranza, U. Sengupta, M. J. Guerrero-Munoz, T. Kiritoshi, V. Neugebauer, G. R. Jackson, R. Kayed, Alzheimer brain-derived tau oligomers propagate pathology from endogenous tau. Sci. Rep. 2, 700 (2012).

30. A. Mudher, M. Colin, S. Dujardin, M. Medina, I. Dewachter, S. M. A. Naini, E.-M. Mandelkow, E. Mandelkow, L. Buée, M. Goedert, What is the evidence that tau pathology spreads through prion-like propagation? Acta. Neuropathol. Commun. 5, 99 (2017).

31. B. Frost, M. I. Diamond, Prion-like mechanisms in neurodegenerative diseases. Nat. Rev. Neurosci. 11, 155–159 (2010).

32. S. K. Sonawane, S. Chinnathambi, Prion-Like Propagation of Post-Translationally Modified Tau in Alzheimer’s Disease: A Hypothesis. J. Mol. Neurosci. 65, 480–490 (2018).

33. S. L. DeVos, B. T. Corjuc, D. H. Oakley, C. K. Nobuhara, R. N. Bannon, A. Chase, C. Commins, J. A. Gonzalez, P. M. Dooley, M. P. Frosch, B. T. Hyman, Synaptic Tau Seeding Precedes Tau Pathology in Human Alzheimer’s Disease Brain. Front. Neurosci. 12, 267 (2018).

34. H. C. Tai, A. Serrano-Pozo, T. Hashimoto, M. P. Frosch, T. L. Spires-Jones, B. T. Hyman, The synaptic accumulation of hyperphosphorylated tau oligomers in Alzheimer disease is associated with dysfunction of the ubiquitin-proteasome system. Am. J. Pathol. 181, 1426–1435 (2012).

35. H. C. Tai, B. Y. Wang, A. Serrano Pozo, M. P. Frosch, T. L. Spires-Jones, B. T. Hyman, Frequent and symmetric deposition of misfolded tau oligomers within presynaptic and postsynaptic terminals in Alzheimer inverted question marks disease. Acta Neuropathol. Commun. 2, 146 (2014).

36. A. Singh, D. Allen, A. Fracassi, B. Tumurbaatar, C. Natarajan, P. Scaduto, R. Woltjer, R. Kayed, A. Limon, B. Krishnan, G. Taglialatela, Functional Integrity of Synapses in the Central Nervous System of Cognitively Intact Individuals with High Alzheimer’s Disease Neuropathology Is Associated with Absence of Synaptic Tau Oligomers. J. Alzheimers Dis. 78, 1661–1678 (2020).

37. M. Colin, S. Dujardin, S. Schraen-Maschke, G. Meno-Tetang, C. Duyckaerts, J. P. Courade, L. Buee, From the prion-like propagation hypothesis to therapeutic strategies of anti-tau immunotherapy. Acta Neuropathol. 139, 3–25 (2020).

38. J.-P. Courade, R. Angers, G. Mairet-Coello, N. Pacico, K. Tyson, D. Lightwood, R. Munro, D. McMillan, R. Griffin, T. Baker, Epitope determines efficacy of therapeutic anti-Tau antibodies in a functional assay with human Alzheimer Tau. Acta Neuropathol. 136, 729–745 (2018).

39. S.-H. Lee, C. E. Le Pichon, O. Adolfsson, V. Gafner, M. Pihlgren, H. Lin, H. Solanoy, R. Brendza, H. Ngu, O. Foreman, R. Chan, J. A. Ernst, D. DiCara, I. Hotzel, K. Srinivasan, D. V. Hansen, J. Atwal, Y. Lu, D. Bumbaca, A. Pfeifer, R. J. Watts, A. Muhs, K. Scearce-Levie, G. Ayalon, Antibody-Mediated Targeting of Tau In Vivo Does Not Require Effector Function and Microglial Engagement. Cell Rep. 16, 1690–1700 (2016).

40. W. R. Galpern, M. Mercken, K. Van Kolen, M. Timmers, K. Haeverans, L. Janssens, G. Triana-Baltzer, H. C. Kolb, T. Jacobs, P. Nandy, T. Malia, H. Sun, L. Van Nueten, P1–052: A single ascending dose study to evaluate the safety, tolerability, pharmacokinetics, and pharmacodynamics of the anti-phospho-tau antibody JNJ-63733657 in healthy subjects. Alzheimers Dement. 15, P252–P253 (2019).

41. N. Rosenqvist, A. A. Asuni, C. R. Andersson, S. Christensen, J. A. Daechsel, J. Egebjerg, J. Falsig, L. Helboe, P. Jul, F. Kartberg, L. O. Pedersen, E. M. Sigurdsson, F. Sotty, K. Skjodt, J. B. Stavenhagen, C. Volbracht, J. T. Pedersen, Highly specific and selective anti-pS396-tau antibody C10.2 targets seeding-competent tau. Alzheimers Dement. 4, 521–534 (2018).

42. L. Collin, B. Bohrmann, U. Göpfert, K. Oroszlan-Szovik, L. Ozmen, F. Grüninger, Neuronal uptake of tau/pS422 antibody and reduced progression of tau pathology in a mouse model of Alzheimer’s disease. Brain 137, 2834–2846 (2014).

43. S. Wegmann, J. Biernat, E. Mandelkow, A current view on Tau protein phosphorylation in Alzheimer’s disease. Curr. Opin. Neurobiol. 69, 131–138 (2021).

44. J. Z. Wang, Y. Y. Xia, I. Grundke-Iqbal, K. Iqbal, Abnormal hyperphosphorylation of tau: sites, regulation, and molecular mechanism of neurofibrillary degeneration. J. Alzheimers Dis. 33 Suppl 1, S123–139 (2013).

45. R. Alam, D. A. Driver, S. Wu, E. Lozano, S. L. Key, J. T. Hole, M. L. Hayashi, J. I. Lu, PRECLINICAL CHARACTERIZATION OF AN ANTIBODY [LY3303560] TARGETING AGGREGATED TAU. Alzheimes Dement. 13, P592–P593 (2017).

46. G. A. Jicha, R. Bowser, I. G. Kazam, P. Davies, Alz-50 and MC-1, a new monoclonal antibody raised to paired helical filaments, recognize conformational epitopes on recombinant tau. J. Neurosci. Res. 48, 128–132 (1997).

47. C. I. Yang, B. N. Tsai, S. J. Huang, T. Y. Wang, H. C. Tai, J. C. Chan, Aggregation of Beta-Amyloid Peptides Proximal to Zwitterionic Lipid Bilayers. Chem. Asian J. 10, 1967–1971 (2015).

48. Z. H. Guo, C. I. Yang, C. I. Ho, S. J. Huang, Y. C. Chen, H. C. Tai, J. C. C. Chan, Fibrillization of beta-Amyloid Peptides via Chemically Modulated Pathway. Chem. Eur. J. 24, 4939–4943 (2018).

49. K. Santacruz, J. Lewis, T. Spires, J. Paulson, L. Kotilinek, M. Ingelsson, A. Guimaraes, M. DeTure, M. Ramsden, E. McGowan, C. Forster, M. Yue, J. Orne, C. Janus, A. Mariash, M. Kuskowski, B. Hyman, M. Hutton, K. H. Ashe, Tau suppression in a neurodegenerative mouse model improves memory function. Science 309, 476–481 (2005).

50. T. Uchihara, Pretangles and neurofibrillary changes: Similarities and differences between AD and CBD based on molecular and morphological evolution. Neuropathology 34, 571–577 (2014).

51. D. J. Selkoe, Alzheimer’s disease is a synaptic failure. Science 298, 789–791 (2002).

52. A. John, P. H. Reddy, Synaptic basis of Alzheimer’s disease: Focus on synaptic amyloid beta, P-tau and mitochondria. Ageing Res. Rev. 65, 101208 (2021).

53. T. L. Spires-Jones, B. T. Hyman, The intersection of amyloid beta and tau at synapses in Alzheimer’s disease. Neuron 82, 756–771 (2014).

54. S. T. DeKosky, S. W. Scheff, Synapse loss in frontal cortex biopsies in Alzheimer’s disease: correlation with cognitive severity. Ann. Neurol. 27, 457–464 (1990).

55. R. D. Terry, E. Masliah, D. P. Salmon, N. Butters, R. DeTeresa, R. Hill, L. A. Hansen, R. Katzman, Physical basis of cognitive alterations in Alzheimer’s disease: synapse loss is the major correlate of cognitive impairment. Ann. Neurol. 30, 572–580 (1991).

56. G. M. Shankar, B. L. Bloodgood, M. Townsend, D. M. Walsh, D. J. Selkoe, B. L. Sabatini, Natural oligomers of the Alzheimer amyloid-β protein induce reversible synapse loss by modulating an NMDA-type glutamate receptor-dependent signaling pathway. J. Neurosci. 27, 2866–2875 (2007).

57. L. M. Ittner, J. Gotz, Amyloid-β and tau--a toxic pas de deux in Alzheimer’s disease. Nat. Rev. Neurosci. 12, 65–72 (2011).

58. E. D. Roberson, K. Scearce-Levie, J. J. Palop, F. Yan, I. H. Cheng, T. Wu, H. Gerstein, G. Q. Yu, L. Mucke, Reducing endogenous tau ameliorates amyloid β-induced deficits in an Alzheimer’s disease mouse model. Science 316, 750–754 (2007).

59. L. M. Ittner, Y. D. Ke, F. Delerue, M. Bi, A. Gladbach, J. van Eersel, H. Wolfing, B. C. Chieng, M. J. Christie, I. A. Napier, A. Eckert, M. Staufenbiel, E. Hardeman, J. Gotz, Dendritic function of tau mediates amyloid-β toxicity in Alzheimer’s disease mouse models. Cell 142, 387–397 (2010).

60. H. Y. Wu, P. C. Kuo, Y. T. Wang, H. T. Lin, A. D. Roe, B. Y. Wang, C. L. Han, B. T. Hyman, Y. J. Chen, H. C. Tai, Beta-Amyloid Induces Pathology-Related Patterns of Tau Hyperphosphorylation at Synaptic Terminals. J. Neuropathol. Exp. Neurol. 77, 814–826 (2018).

61. T. Petrozziello, E. A. Bordt, A. N. Mills, S. E. Kim, E. Sapp, B. A. Devlin, A. A. Obeng-Marnu, S. M. K. Farhan, A. C. Amaral, S. Dujardin, P. M. Dooley, C. Henstridge, D. H. Oakley, A. Neueder, B. T. Hyman, T. L. Spires-Jones, S. D. Bilbo, K. Vakili, M. E. Cudkowicz, J. D. Berry, M. DiFiglia, M. C. Silva, S. J. Haggarty, G. Sadri-Vakili, Targeting Tau Mitigates Mitochondrial Fragmentation and Oxidative Stress in Amyotrophic Lateral Sclerosis. Mol. Neurobiol. 59, 683–702 (2022).

62. T. Smolek, A. Madari, J. Farbakova, O. Kandrac, S. Jadhav, M. Cente, V. Brezovakova, M. Novak, N. Zilka, Tau hyperphosphorylation in synaptosomes and neuroinflammation are associated with canine cognitive impairment. J. Comp. Neurol. 524, 874–895 (2016).

63. S. Sokolow, K. M. Henkins, T. Bilousova, B. Gonzalez, H. V. Vinters, C. A. Miller, L. Cornwell, W. W. Poon, K. H. Gylys, Pre-synaptic C-terminal truncated tau is released from cortical synapses in Alzheimer’s disease. J. Neurochem. 133, 368–379 (2015).

64. T. E. Tracy, J. Madero-Perez, D. L. Swaney, T. S. Chang, M. Moritz, C. Konrad, M. E. Ward, E. Stevenson, R. Huttenhain, G. Kauwe, M. Mercedes, L. Sweetland-Martin, X. Chen, S. A. Mok, M. Y. Wong, M. Telpoukhovskaia, S. W. Min, C. Wang, P. D. Sohn, J. Martin, Y. Zhou, W. Luo, J. Q. Trojanowski, V. M. Y. Lee, S. Gong, G. Manfredi, G. Coppola, N. J. Krogan, D. H. Geschwind, L. Gan, Tau interactome maps synaptic and mitochondrial processes associated with neurodegeneration. Cell 185, 712–728 e714 (2022).

65. Y. Wang, V. Balaji, S. Kaniyappan, L. Kruger, S. Irsen, K. Tepper, R. Chandupatla, W. Maetzler, A. Schneider, E. Mandelkow, E. M. Mandelkow, The release and trans-synaptic transmission of Tau via exosomes. Mol. Neurodegener. 12, 5 (2017).

66. A. W. P. Fitzpatrick, B. Falcon, S. He, A. G. Murzin, G. Murshudov, H. J. Garringer, R. A. Crowther, B. Ghetti, M. Goedert, S. H. W. Scheres, Cryo-EM structures of tau filaments from Alzheimer’s disease. Nature 547, 185–190 (2017).

67. T. K. Karikari, D. A. Nagel, A. Grainger, C. Clarke-Bland, J. Crowe, E. J. Hill, K. G. Moffat, Distinct Conformations, Aggregation and Cellular Internalization of Different Tau Strains. Front. Cell. Neurosci. 13, 296 (2019).

68. R. Abskharon, P. M. Seidler, M. R. Sawaya, D. Cascio, T. P. Yang, S. Philipp, C. K. Williams, K. L. Newell, B. Ghetti, M. A. DeTure, D. W. Dickson, H. V. Vinters, P. L. Felgner, R. Nakajima, C. G. Glabe, D. S. Eisenberg, Crystal structure of a conformational antibody that binds tau oligomers and inhibits pathological seeding by extracts from donors with Alzheimer’s disease. J. Biol. Chem. 295, 10662–10676 (2020).

69. A. Gómez-Ramos, M. Díaz-Hernández, R. Cuadros, F. Hernández, J. Avila, Extracellular tau is toxic to neuronal cells. FEBS Lett. 580, 4842–4850 (2006).

70. C. A. Lasagna-Reeves, D. L. Castillo-Carranza, U. Sengupta, J. Sarmiento, J. Troncoso, G. R. Jackson, R. Kayed, Identification of oligomers at early stages of tau aggregation in Alzheimer’s disease. FASEB J. 26, 1946–1959 (2012).

71. D. L. Castillo-Carranza, U. Sengupta, M. J. Guerrero-Munoz, C. A. Lasagna-Reeves, J. E. Gerson, G. Singh, D. M. Estes, A. D. Barrett, K. T. Dineley, G. R. Jackson, R. Kayed, Passive immunization with Tau oligomer monoclonal antibody reverses tauopathy phenotypes without affecting hyperphosphorylated neurofibrillary tangles. J. Neurosci. 34, 4260–4272 (2014).

72. K. R. Patterson, C. Remmers, Y. Fu, S. Brooker, N. M. Kanaan, L. Vana, S. Ward, J. F. Reyes, K. Philibert, M. J. Glucksman, L. I. Binder, Characterization of prefibrillar Tau oligomers in vitro and in Alzheimer disease. J. Biol. Chem. 286, 23063–23076 (2011).

73. B. Wolozin, P. Davies, Alzheimer-related neuronal protein A68: specificity and distribution. Ann. Neurol. 22, 521–526 (1987).

74. K. Pang, R. Jiang, W. Zhang, Z. Yang, L. L. Li, M. Shimozawa, S. Tambaro, J. Mayer, B. Zhang, M. Li, J. Wang, H. Liu, A. Yang, X. Chen, J. Liu, B. Winblad, H. Han, T. Jiang, W. Wang, P. Nilsson, W. Guo, B. Lu, An App knock-in rat model for Alzheimer’s disease exhibiting Abeta and tau pathologies, neuronal death and cognitive impairments. Cell Res. 32, 157–175 (2022).

75. L. A. Gomes, V. Uytterhoeven, D. Lopez-Sanmartin, S. O. Tomé, T. Tousseyn, R. Vandenberghe, M. Vandenbulcke, C. A. von Arnim, P. Verstreken, D. R. Thal, Maturation of neuronal AD-tau pathology involves site-specific phosphorylation of cytoplasmic and synaptic tau preceding conformational change and fibril formation. Acta Neuropathol. 141, 173–192 (2021).

76. D. J. Koss, G. Jones, A. Cranston, H. Gardner, N. M. Kanaan, B. Platt, Soluble pre-fibrillar tau and β-amyloid species emerge in early human Alzheimer’s disease and track disease progression and cognitive decline. Acta Neuropathol. 132, 875–895 (2016).

77. D. J. Koss, M. Dubini, H. Buchanan, C. Hull, B. Platt, Distinctive temporal profiles of detergent-soluble and -insoluble tau and Abeta species in human Alzheimer’s disease. Brain Res. 1699, 121–134 (2018).

78. N. Sahara, Y. Ren, S. Ward, L. I. Binder, T. Suhara, M. Higuchi, Tau oligomers as potential targets for early diagnosis of tauopathy. J. Alzheimers Dis. 40 Suppl 1, S91–96 (2014).

79. Z. He, J. D. McBride, H. Xu, L. Changolkar, S.-j. Kim, B. Zhang, S. Narasimhan, G. S. Gibbons, J. L. Guo, M. Kozak, Transmission of tauopathy strains is independent of their isoform composition. Nature Comm. 11, 1–18 (2020).

80. S. Narasimhan, J. L. Guo, L. Changolkar, A. Stieber, J. D. McBride, L. V. Silva, Z. He, B. Zhang, R. J. Gathagan, J. Q. Trojanowski, Pathological tau strains from human brains recapitulate the diversity of tauopathies in nontransgenic mouse brain. J. Neurosci. 37, 11406–11423 (2017).

81. B. Falcon, W. Zhang, A. G. Murzin, G. Murshudov, H. J. Garringer, R. Vidal, R. A. Crowther, B. Ghetti, S. H. Scheres, M. Goedert, Structures of filaments from Pick’s disease reveal a novel tau protein fold. Nature 561, 137–140 (2018).

82. B. Falcon, W. Zhang, M. Schweighauser, A. G. Murzin, R. Vidal, H. J. Garringer, B. Ghetti, S. H. Scheres, M. Goedert, Tau filaments from multiple cases of sporadic and inherited Alzheimer’s disease adopt a common fold. Acta Neuropathol. 136, 699–708 (2018).

83. W. Zhang, A. Tarutani, K. L. Newell, A. G. Murzin, T. Matsubara, B. Falcon, R. Vidal, H. J. Garringer, Y. Shi, T. Ikeuchi, Novel tau filament fold in corticobasal degeneration. Nature 580, 283–287 (2020).

84. C. d’Abramo, C. M. Acker, H. T. Jimenez, P. Davies, Tau passive immunotherapy in mutant P301L mice: antibody affinity versus specificity. PLoS One 8, e62402 (2013).

85. S. Maeda, N. Sahara, Y. Saito, M. Murayama, Y. Yoshiike, H. Kim, T. Miyasaka, S. Murayama, A. Ikai, A. Takashima, Granular tau oligomers as intermediates of tau filaments. Biochemistry 46, 3856–3861 (2007).

86. S. Schroeder, A. Joly-Amado, A. Soliman, U. Sengupta, R. Kayed, M. N. Gordon, D. Morgan, Oligomeric tau-targeted immunotherapy in Tg4510 mice. Alzheimers Res. Ther. 9, 46 (2017).

87. M. Maruyama, H. Shimada, T. Suhara, H. Shinotoh, B. Ji, J. Maeda, M. R. Zhang, J. Q. Trojanowski, V. M. Lee, M. Ono, K. Masamoto, H. Takano, N. Sahara, N. Iwata, N. Okamura, S. Furumoto, Y. Kudo, Q. Chang, T. C. Saido, A. Takashima, J. Lewis, M. K. Jang, I. Aoki, H. Ito, M. Higuchi, Imaging of tau pathology in a tauopathy mouse model and in Alzheimer patients compared to normal controls. Neuron 79, 1094–1108 (2013).

88. T. G. Beach, C. H. Adler, L. I. Sue, G. Serrano, H. A. Shill, D. G. Walker, L. Lue, A. E. Roher, B. N. Dugger, C. Maarouf, A. C. Birdsill, A. Intorcia, M. Saxon-Labelle, J. Pullen, A. Scroggins, J. Filon, S. Scott, B. Hoffman, A. Garcia, J. N. Caviness, J. G. Hentz, E. Driver-Dunckley, S. A. Jacobson, K. J. Davis, C. M. Belden, K. E. Long, M. Malek-Ahmadi, J. J. Powell, L. D. Gale, L. R. Nicholson, R. J. Caselli, B. K. Woodruff, S. Z. Rapscak, G. L. Ahern, J. Shi, A. D. Burke, E. M. Reiman, M. N. Sabbagh, Arizona Study of Aging and Neurodegenerative Disorders and Brain and Body Donation Program. Neuropathology 35, 354–389 (2015).

